# Computational screening of potential AT1R inhibitors from *Nigella sativa* for diabetic-hypertensive therapy

**DOI:** 10.1101/2024.08.30.610463

**Authors:** Tonima Rahman Tuli, Mijan Mia, SK Nazmul Ulla, Md. Shamim Gazi, Md. Nazmul Hasan, Partha Biswas, Md. Mohaimenul Islam Tareq

**Affiliations:** Biotechnology and Genetic Engineering Discipline, Life Science School, Khulna University, Khulna, Bangladesh; Department of Genetic Engineering and Biotechnology, Jashore University of Science and Technology, Jashore 7408, Bangladesh

**Keywords:** Angiotensin II, Angiotensin-II Type1 Receptor (AT1R), Molecular Simulation, Diabetes, Hypertension

## Abstract

Hypertension, frequently coexists with Type 2 diabetes mellitus, causes due to Angiotensin II. Inhibiting Angiotensin II by targeting AT1R can potentially improve hypertension-related complications in patients with T2DM. In this study, we perform an in-silico screening based on molecular docking and molecular dynamic of 305 compounds of Nigella sativa against Angiotensin-II Type1 Receptor (AT1R). Molecular docking studies were conducted to investigate the binding affinities of the selected ligands with the target receptor and compared against the Losartan (control drug). Top 5 ligands were selected based on binding affinity. Subsequently, we conducted an ADMET (Absorption, Distribution, Metabolism, Excretion, and Toxicity) profiling to assess their pharmacokinetic and safety profiles. Our findings revealed two promising ligands Beta-amyrin (CID-73145) and Taraxerol (CID-92097), which exhibited strong binding affinities and favorable pharmacological profiles with no signs of acute toxicity. The molecular dynamic simulations (MD) including RMSD, RMSF, and MMGBSA binding free energy results showed that selected two ligands bind to AT1R more proficiently with good stability over 100 ns. The computational study serves as a foundational basis for subsequent laboratory and clinical research aimed at identifying selective, potent, and less toxic diabetic-hypertensive therapy targeting AT1R.

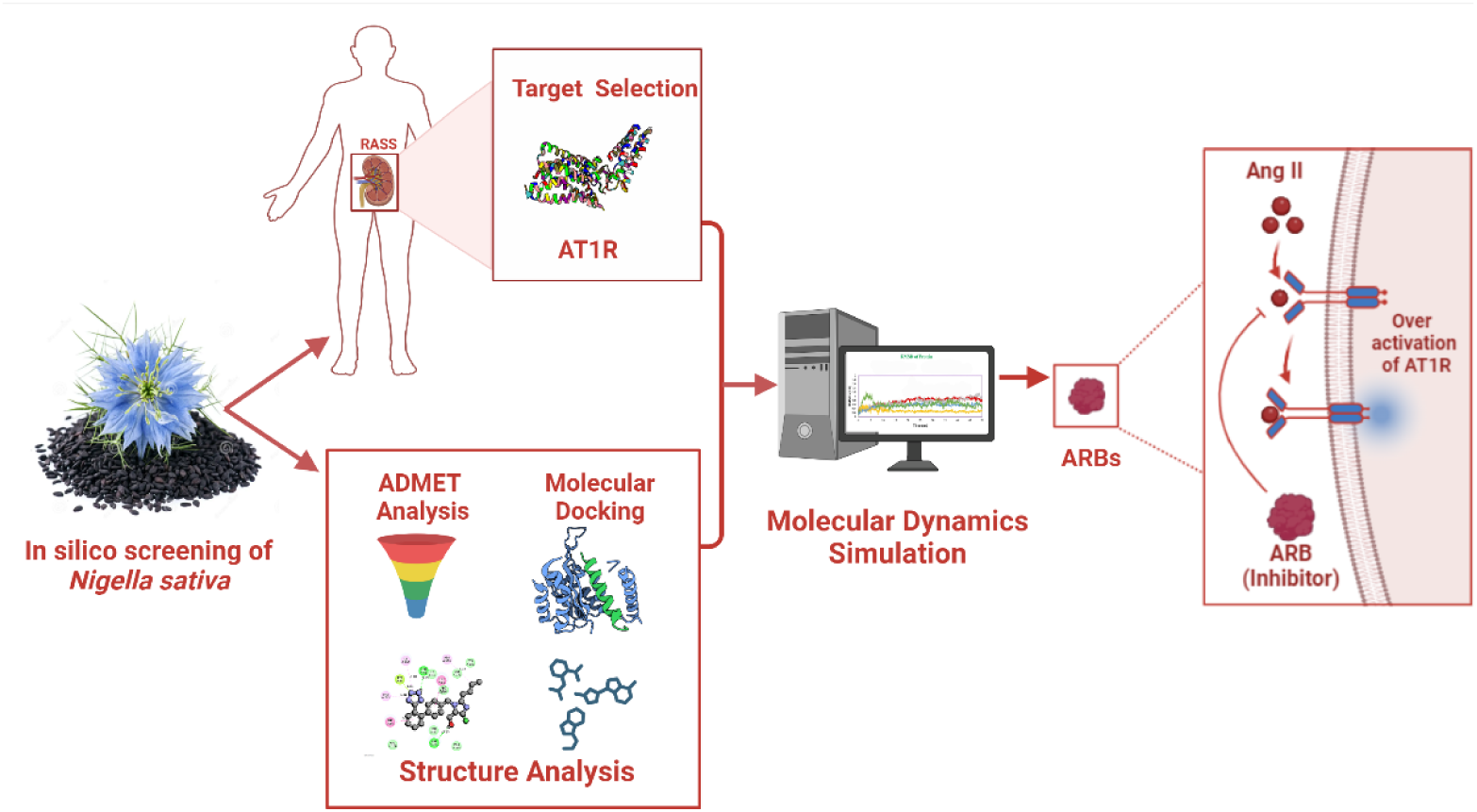

## 1. Introduction

Hypertension is a crucial factor that causes cardiovascular disease (CVD) and a fourth major cause of premature death ^1^. Approximately 17 million deaths each year result from cardiovascular diseases, with hypertension alone accounting for 9.4 million of these fatalities^2^. As stated by the World Health Organization (WHO), currently 1.13 billion individuals are enduring hypertension worldwide. By 2025, The rise in the prevalence of hypertension among adults is projected to be 60%, reaching a total of 1.56 billion ^3^. Tabassum and Ahmad (2011) stated hypertension as a CVD having a systolic blood pressure (SBP) and DBP of more than 140/90 mmHg ^4^. Hypertension occurs due to the activity of the Ang-II hormone. It is one of the root causes of the onset of many other diseases, among them type 2 diabetes mellitus (T2DM) is prevalent.

Hypertension is a common phenomenon in T2DM^5^. The strong correlation between diabetes and hypertension has been acknowledged for decades ^6^. The premature mortality rate in diabetic-hypertensive patients is over 50% and the leading cause is hypertension. An altered glucose metabolism is responsible for diabetes in hypertension. In high blood pressure, the presence of excess Ang-II often mutates the bio functionality of blood-glucose maintaining hormone insulin that governs the Insulin resistance. This phenomenon is one of the leading causes of diabetes, is frequently observed in individuals with hypertension. Consequently there is a clear association between the levels of plasma insulin concentrations and blood pressure (BP), that emerges the diabetes complication ^7^. Therefore, effective hypertension management can simultaneously diminish the progression of cardiovascular and diabetes complications in patients with both conditions. ^8^.

Angiotensin-II is an octapeptide vasoconstrictor hormone produced by the enzymatic cascade reaction of renin angiotensin–aldosterone system (RAAS) ^9^. This hormone administrates the CV homeostasis by ensuring an equilibrium between blood volume and vascular resistance ^10^. There are two sub-families of Ang-II namely angiotensin II type 1 receptor (AT1R) and type 2 (AT2R), that are two well-defined G-protein-coupled receptors (GPCR), respond to the angiotensin-II ^11^. But AT1 receptor demonstrates a maximum tendency to exert critical function in vasoconstriction which comprises hypertension or increased blood pressure. Thereby, AT1R regulates the majority of cardiovascular activities alongside regulating and sustaining oxidative stress, aldosterone release, vascular constriction, kidney sodium retention, vasopressin secretion, and other factors ^12^. Angiotensin-II is released by RAAS due to multiple factors and binds to the AT1R. But whenever Ang-II over-activates the AT1R, in response the vascular system constricts the blood vessels, elevates the blood pressure, and eventually results in hypertension ^13^. Hence, scientists are strategically targeting the AT1R than the AT2R for evolving promising treatment and anti-hypertension drugs and other cardiovascular ailments ^14^.

Angiotensin-II type 1 receptor blockers (ARBs) are novel drugs with dual potential as anti-hypertensive agents and for addressing T2DM. These medications function by obstructing the Ang-II from attaching to the AT1R. Thereby, ARBs balance the level of Ang II in the body by regulating the salt and water concentration. This leads to vasodilation of the blood vessels and maintenance of blood pressure through the inhibition of AT1R ^15^. Moreover, ARBs impede the activity of RAAS which is a key player in the vascular damage, responsible for comprising numerous severe complications associated with diabetes ^16^.

Angiotensin-II hormone inhibits the insulin signaling pathways, which activates the superoxide and reactive oxygen species (ROS) or nitric oxide production ^17^. Interrupting the RAAS by ARBs offer lowering the oxidative stress and enhancing nitric oxide production ^18^. These in turn, can result in enhanced insulin signaling for potential improvements in diabetes. Evidence suggests that RAAS has a potential impact on pancreatic β-cell function and insulin-secreting capacity altering the islet perfusion and insulin secretion ^19^. Therefore, inhibiting RAAS maintains pancreatic β-cell function and insulin secretion capability, resulting in enhanced insulin secretion and improved glucose metabolism in diabetes ^20^. ARBs can contribute to improved insulin delivery, promoting glucose absorption and metabolism in insulin-sensitive tissues by enhanced muscle blood flow associated with the attenuation of the RAAS ^21, 22^. Moreover, increased vascular sensitivity can enhance insulin delivery, subsequently contributing to more effective glucose utilization in tissues interacting with peroxisome proliferator-activated receptor γ. This suggest a mechanism for improving insulin sensitivity independent of angiotensin II receptors [24][26]. All these mechanisms suggest the improvement of diabetes conditions in diabetic-hypertensive patients by consuming ARBs. Currently, eight ARBs– Losartan, Telmisartan, Olmesartan, Valsartan Azilsartan, Eprosartan, Candesartan, and Irbesartan– are available for clinical use ^14^. However, due to side effects, less long-term efficacy, and being expensive, the development of novel drugs is the urgency of the time. Therefore, considering the pervious study and evidence, it is prudent that the phytochemicals can be used as potent and efficacious angiotensin receptor blockers to provide cardio protection and improve diabetes.

Nigella sativa (NS), also known as black seed or black cumin, is an annual dicotyledonous herb and a member of the Ranunculaceae family, is used as an indigenous natural medicine all over the world ^26,27^. NS is reported as a natural remedy that holds several therapeutic uses, along with diabetes, tumors, hypercholesterolemia, hypertension, inflammation, and gastrointestinal disorders ^28^ ^29,30^. Besides, its active components are clinically proven to exert antioxidant, hypotensive, calcium channel blocking, and diuretic properties, potentially playing a role in lowering blood pressure ^31^. The diuretic functionality of NS, marked by the elevated elimination of Na^+^, K^+,^ Cl^−^, and urea, implies its potentiality to lower blood pressure through a reduction in body electrolytes and water, subsequently decreasing blood volume and cardiac output, key factors in blood pressure regulation. NS has been reported to have the capability to lower blood pressure through its influence on calcium (Ca^2+)^ ion channels. NS blocks the Ca^2+^ channel, and inhibit its release from the sarcoplasmic reticulum, a decrease in Ca2+ entry into smooth muscle cells in blood vessels ultimately results in increased vasorelaxation ^32^ This study suggests the bimodal therapeutic potential of the components of NS as ARB using employing in silico techniques and cheminformatic analysis to examine the therapeutic mode of NS in the management of hypertension, and thus improve diabetes ^33^.

Computer-aided drug designing (CADD) incorporates, molecular docking, ADME analysis, toxicity profiling and molecular dynamics simulations. These methods predict how drugs bind to target proteins, offering insights into H-bond, hydrophobic, Van Der Waals and electrostatic interactions. This information helps employ structure-based drug design to scrutinize binding affinities, drug-like properties, bioavailability, and ADMET properties. Molecular dynamics simulations then enlightened the underlying interactions dynamically, mimicking cellular conditions for greater accuracy in predicting stability, flexibility and thermodynamic characteristics of the target protein and lead^34^ ^35^. In this present study, we have assessed the blocking potential of the phytochemicals of NS against Ang-II-the protein responsible for various cardiac diseases and compared the activities with known angiotensin ARBs.

## 2. Experimental section

### 2.1. Validation and active site prediction

The 3D X-ray crystal structure of our target protein AT1R (PDB ID: 4ZUD), was acquired from RCSB PDB. The protein consisted of 410 amino acids, was in complex with inverse agonist Olmesartan with a low free R-value (0.234) and the structural resolution was (2.80Å) ^36^. For validation of protein, Firstly We utilized the PROCHECK server to generate a Ramachandran plot, which provided an accurate representation of the residues within allowed and disallowed regions ^37^. Furthermore we used the ProSa web server to generate the Z score of the structure as the score shows overall protein model quality and estimates the deviation of the total energy of the structure^38^. The active sites of the receptor-protein AT1R were unveiled using the CASTp server (Computed Atlas of Surface Topography of Proteins) web tool. It also allows the demonstration of the volume and area of the binding pocket from crystal structures ^39^.

### 2.2 Optimization of protein AT1R

In this study, the protein was optimized for primary structure preparation to execute docking studies using UCSF Chimera.^40^ The multistage processes followed the elimination of co-crystalized water molecules, nonpolar hydrogens, small molecules, nonstandard residues, ion pairs and the addition of polar hydrogens to upgrade the target structure ^41^.

### 2.3. Ligand reclamation and preparation

This study investigated 305 compounds of *Nigella sativa* for molecular docking against AT1R, from the Indian Medicinal Plants, Phytochemistry, And Therapeutics (IMPPAT), which is a manually compiled database. The entitled phytochemicals were retrieved and assembled in 3D SDF format from PubChem for virtual screening. Furthermore, “Losartan” an FDA-approved angiotensin receptor blocker (ARB) drug served as a positive control ^42^ ^43^. In PyRx program, using Open Babel, the energy of the ligands was diminished to become relaxed and transformed into a PDBQT file ^44^.

### 2.4. Virtual screening and molecular docking

The molecular docking study was executed with the assistance of AutoDock Vina Wizard using the PyRx software to discover the most suitable binding configurations of the protein and ligand. This is the key component for anticipating virtual screening in structural biology, especially for CADD. We screened 305 compounds of *Nigella sativa* at the prior level considering as ligands via molecular docking against the target protein ^45^. Before conducting molecular docking, AT1R was transformed into ‘pdbqt’ format, and a grid box center (x, y, z) = 20.1780, 51.1697, 56.2881, and dimensions (x, y, z) = 51.4161, 51.1697, 56.2881 was developed to encompass the residues revealed in the binding site cavity which allows the free movement of the ligands. We performed docking of reported ligands against the target protein to dock with all possible binding sites of the protein. The docking score was represented as a negative score in kcal/mol, where the highest negative docking score indicates the strongest binding energy. The derivatives with the lowest scores were then submitted to ADMET and drug-likeness analysis.

### 2.5 Drug-likeness, ADME study and Toxicity analysis

The ADMET (absorption, distribution, metabolism, excretion and toxicity) analysis of top ligands based on their binding score was performed using web server SwissADME ^46^. The ADMET profiling was carried out for top compounds based on their binding energy. The canonical SMILES of the compounds were uncovered from the PubChem and submitted to the SwissADME to assess their drug-likeness properties and document their PK values. The online server provided reliable data for assessing physicochemical properties, such as pharmacokinetics, water solubility, lipophilicity, toxicity, and drug-likeness. The evaluation of drug-likeness for the compounds was conducted using ‘Lipinski’s Rule of Five’, Ghose, Veber, and Egans’ selection criteria ^47^. The properties of the compounds having adverse effects on human health were predicted at various levels using the Protox-3.0 server. This study utilized the web server to determine acute toxicity, mutagenicity hepatotoxicity, cytotoxicity, carcinogenicity and immunotoxicity targets of the chosen compounds ^48^.

### 2.6 Post docking analysis of AT1R complex

The compounds with the best docking score as well as good ADMET profiles were further investigated based on their 2D and 3D interactions. We used the PyRx 0.8 and Discovery Studio Visualizer to confirm the best pose for each of the compounds. Biovia software was used to visualize the 2D and 3D molecular interactions between protein(AT1R) and ligands ^49^. This tool showcases a spectrum of interactions within the receptor-ligand docked complex, offering detailed insights into their dynamic interplay.

### 2.7. Molecular Dynamic Simulation

Molecular dynamics simulations lasting 100 ns were employed to assess the interaction flexibility of specific compounds with the binding site residues of AT1R. Compound selection for molecular dynamics simulations was based on superior binding affinities and favorable pharmacokinetic properties identified through molecular docking. The molecular dynamics simulation of the protein complex structures was carried out by using the “Desmond v3.6 Program” in Schrodinger (https://www.schrodinger.com/) (Paid version) had been used within a Linux framework for performing molecular dynamic simulations evaluating various protein-ligand complex structures^50,51^. Employing this framework, the predefined TIP3P aqueous strategy was utilized to create a predetermined volume with an orthorhombic periodic bounding box shape, divided into intervals of 10 Å. Additionally, recommended ions, such as 0.15 M salt (Na+ and Cl-), were selected and dispersed randomly throughout the chemical solvent environment to neutralize electric charge within the structure. After building its solvency protein structures containing agonist combinations, the system’s framework subsequently reduced as well as comfortable utilizing the protocol carried out applying force field constants OPLS3e included inside the Desmond package. Each Isothermal-Isobaric ensemble (NPT) assembly maintained an overall Nose-Hoover temperature combination, ensuring isotropic technique, at approximately 300 K and one atmospheric pressure (1.01325 bar). Following this, there were 50 PS grabbing pauses with an efficiency of 1.2 kcal/mol. This fidelity with its MD simulation was assessed throughout its whole simulation applying the Simulations Interaction Diagram (SID) from the Desmond modules from this Schrödinger suite^51,52^.

### 2.8 Calculation of binding free energy using MMGBSA

To assess the free energies of the complexes throughout the 100 ns simulation, we utilized the thermal mmgbsa Python package to calculate MMGBSA ^53^. The Desmond MD trajectory was divided into 20 individual frame snapshots, and each snapshot underwent MMGBSA analysis, with subsequent separation of the ligand from the receptor. The docked poses underwent local optimization to refine their conformations, followed by computation of the complex’s energies using the OPLS_2005 force field (Schrodinger Release 2023-1). Subsequently, the binding free energy (DG (bind)) values were calculated using the equation:

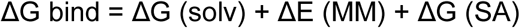

Where DG (solv) =Difference in the GBSA solvation energy of the complex and the sum of the solvation energies for the ligand and unliganded protein, DE(MM)= Energy difference between the complex structure and the sum of the energies of the protein with and without ligand, and DG(SA)= Difference in the surface area energy for the complex and the sum of the surface area energies for the ligand of an uncomplexed protein.

## 3. Results

### 3.1 Structure validation and optimization of AT1R

In order to assess the accuracy of the protein structure, we wanted to create a Ramachandran plot which serves as works as a verification tool indicating the geometries of the protein don’t belong to an electrostatically unfavored area. The Ramachandran plot of the protein AT1R structure was obtained using the Procheck web server and the plot depicts that there were 92.4% atoms in the most favored region and 7.6% atoms in extra allowed regions. For checking the overall protein quality, we checked the Z score which was -4.8 (Figure 1). The result was satisfactory and therefore we finalized subjecting the protein to molecular docking. After extensive optimization of the target protein, we were now ready for molecular docking studies.

**Figure 1:**
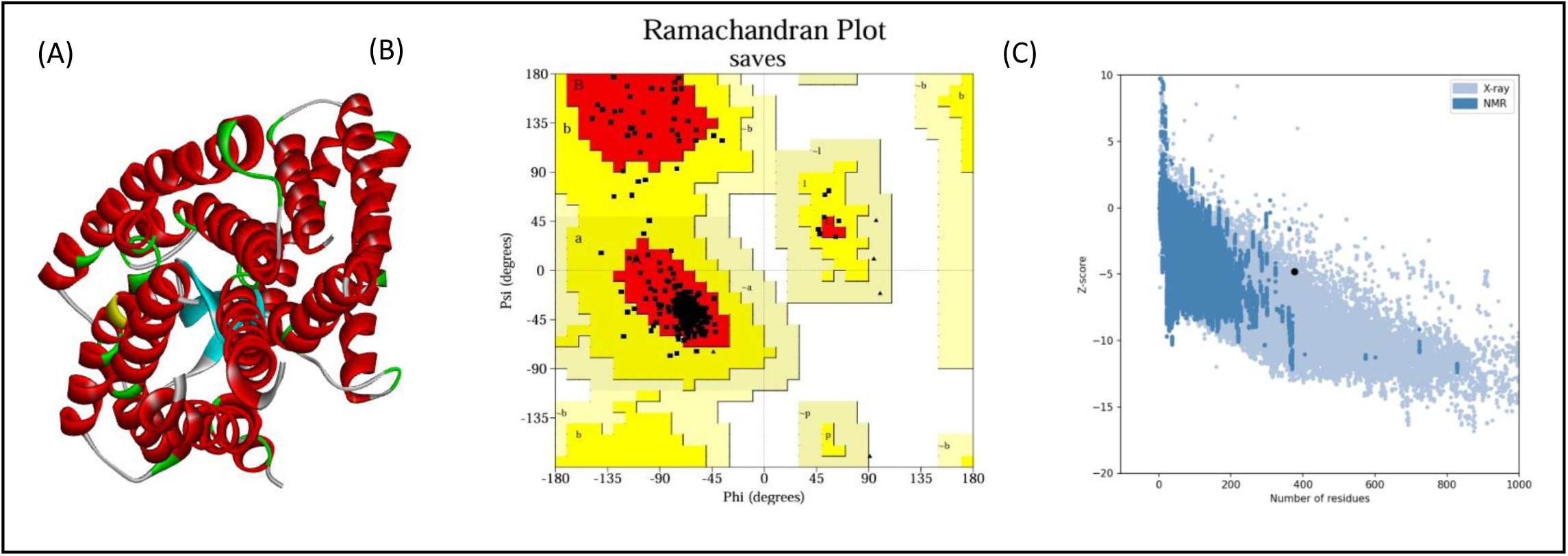
(A) The optimized structure of AT1R without any water molecules, heteroatoms, ions and native ligands (B) Ramachandran plot of AT1R showcasing majority of residues in allowed region (C) Z score of the target protein within an acceptable range.

### 3.2. Phytochemical retrieval and preparation

The IMPPAT database was initially searched to retrieve the available compounds of the target plant ^54^. The database documented 305 compounds from the Nigella sativa (black cumin) plant listed in Supporting Information Table 1. The phytochemical compounds found in the black cumin plant were downloaded and saved in a 2D (SDF) file format. The compounds were prepared and optimized during the ligand preparation steps and evaluation then transformed into the pdbqt file format for additional evaluation.

**Table 1:**
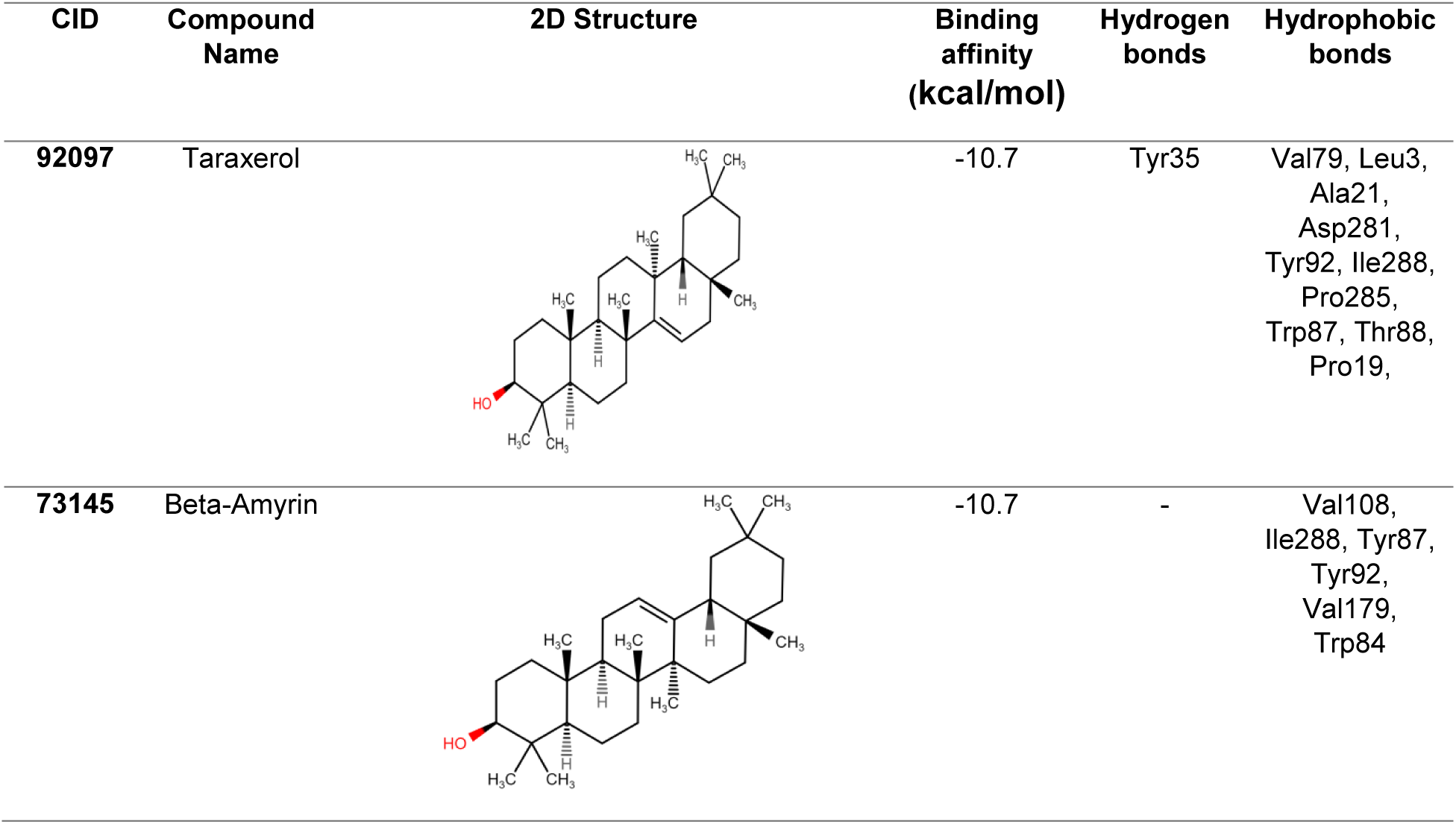
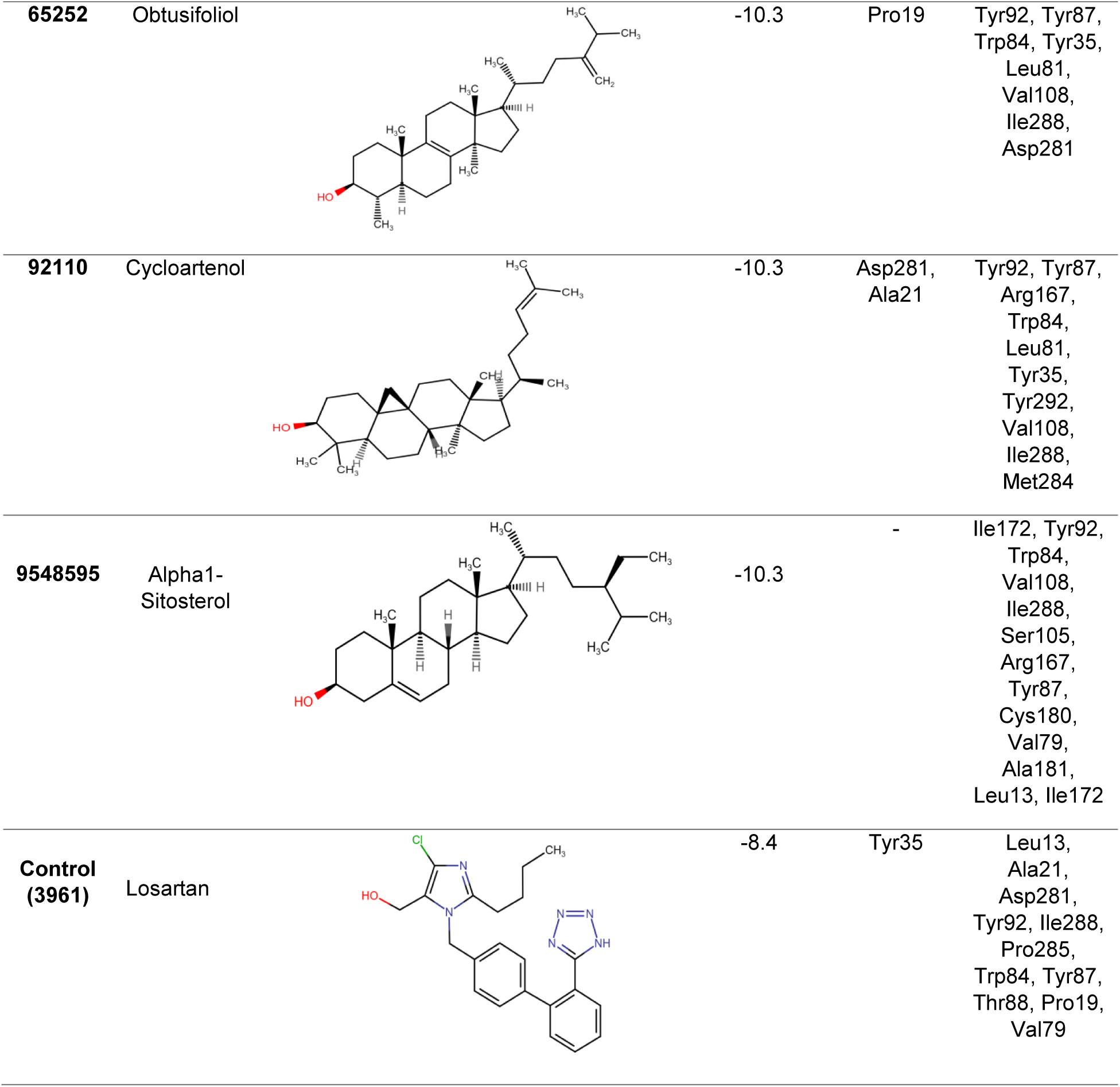
List of selected compounds and control and their CID, compound name, 2D structure, binding affinity, residues of hydrogen bonds and hydrophobic bonds.

| CID | Compound Name | 2D Structure | Binding affinity (kcal/mol) | Hydrogen bonds | Hydrophobic bonds |
| --- | --- | --- | --- | --- | --- |
| 92097 | Taraxerol |  | -10.7 | Tyr35 | Val79, Leu3, Ala21, Asp281, Tyr92, Ile288, Pro285, Trp87, Thr88, Pro19, |
| 73145 | Beta-Amyrin |  | -10.7 | - | Val108, Ile288, Tyr87, Tyr92, Val179, Trp84 |
| 65252 | Obtusifoliol |  | -10.3 | Pro19 | Tyr92, Tyr87, Trp84, Tyr35, Leu81, Val108, Ile288, Asp281 |
| 92110 | Cycloartenol |  | -10.3 | Asp281, Ala21 | Tyr92, Tyr87, Arg167, Trp84, Leu81, Tyr35, Tyr292, Val108, Ile288, Met284 |
| 9548595 | Alpha1-Sitosterol |  | -10.3 | - | Ile172, Tyr92, Trp84, Val108, Ile288, Ser105, Arg167, Tyr87, Cys180, Val79, Ala181, Leu13, Ile172 |
| Control (3961) | Losartan |  | -8.4 | Tyr35 | Leu13, Ala21, Asp281, Tyr92, Ile288, Pro285, Trp84, Tyr87, Thr88, Pro19, Val79 |

### 3.3 Study of the catalytic sites of AT1R

In the case of Angiotensin II type-1 receptor (AT1R), the active sites of the protein were recognized using the CASTp server. The accessible surface area of the active sites was 1820.859 Å. There are 42 active site residues of Angiotensin type-1 receptor-ll (AT1R) as we can visualize from Figure 2.

**Figure 2:**
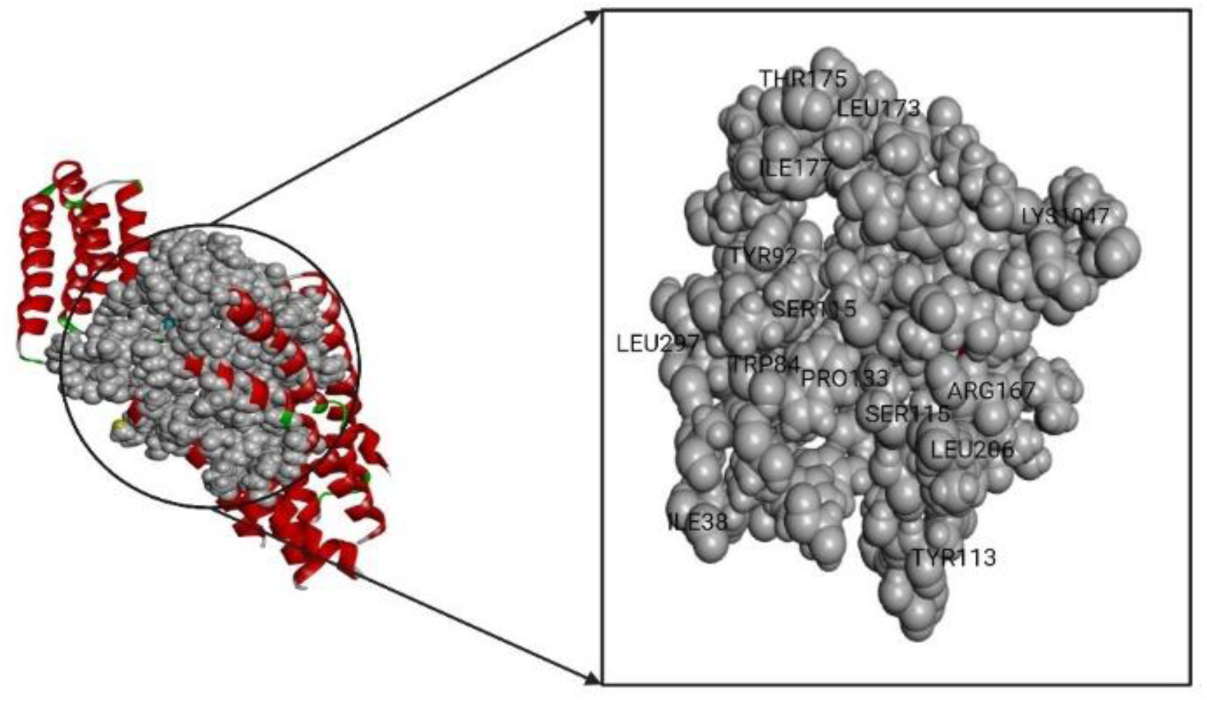
The structure of Angiotensin type-1 receptor-ll (AT1R) where the active sites are shown in ball-shaped formation

### 3.4 Molecular docking-based screening

Computational molecular docking study determines the prime intermolecular arrangements between a protein and a drug candidate. At the very beginning, molecular docking was conducted with PyRx tools AutoDock Vina wizard, between ligands from NS and the AT1R to determine the binding energies. The docking result explicated that the binding affinities were ranged between −3.1 and −10.7 and kcal/mol. Top 5 out of 305 phytochemicals were chosen, based on the binding affinity which has a better binding affinity than control drug Losartan Table 1. The binding affinities of all ligands are provided in Supporting Information Table S1.

### 3.5 Drug likeness and ADMET properties of the selected compounds

The viability assessment of drug discovery involves analyzing both toxicity and drug-likeness through ADMET analysis. The primary concern of ADME is the movement of medications in and out of the body with respect to their dosage and length of time. ^55^. Various physiochemical, pharmacokinetic and drug-like properties were retrieved from SwissADME and pkCSM online tool and documented in Table 2. The analysis assures no violation of Lipinski’s rule of five (RO5). Five compounds including their names, all showed H-bond donor of less than 5, H-bond acceptor of less than 10, molecular weight of less than 500, ClogP greater than 5 which corresponded with the Lipinski and Ghose rule ^56^. The rapid progress of in silico toxicity is emerging as a critical platform for predicting the harmful effects of chemicals on humans, animals, plants, and the environment. ^57^. Hepatoxicity, carcinogenicity, mutagenicity, immunogenicity, cytotoxicity, Ames toxicity and cardiotoxicity factor hERG I/II inhibitor (cardiac toxicity) were analyzed of the selected compounds and the summary is shown in Table 3.

**Table 2:** List of all the pharmacological properties-molecular weight, hydrogen bond donor, hydrogen bond acceptor, rotatable bonds, WLogP (E SOL), gastrointestinal absorption (%), bioavailability, blood-brain barrier permeability, and violations of the rule of five the selected compounds.

| Compound Name | MW | HBD | HBA | RB | WLogP | (E SOL) | GIA (%) | BA | BBB P. | V. of RO5 |
| --- | --- | --- | --- | --- | --- | --- | --- | --- | --- | --- |
| Taraxerol | 426.72 | 1 | 1 | 4 | 8.34 | -6.31 | 94.036 | 0.55 | No | 1 |
| Beta-Amyrin | 426.72 | 1 | 1 | 0 | 8.17 | -6.531 | 93.733 | 0.55 | No | 1 |
| Obtusifoliol | 426.72 | 1 | 1 | 0 | 8.17 | -7.284 | 93.264 | 0.55 | No | 1 |
| Cycloartenol | 426.72 | 1 | 1 | 5 | 8.34 | -5.762 | 95.248 | 0.55 | No | 1 |
| Alpha1-Sitosterol | 426.72 | 1 | 1 | 4 | 8.17 | -6.662 | 94.878 | 0.55 | No | 1 |
| Losartan-Control | 422.91 | 2 | 5 | 8 | 4.11 | -2.915 | 79.338 | 0.56 | No | 0 |

**Table 3:**
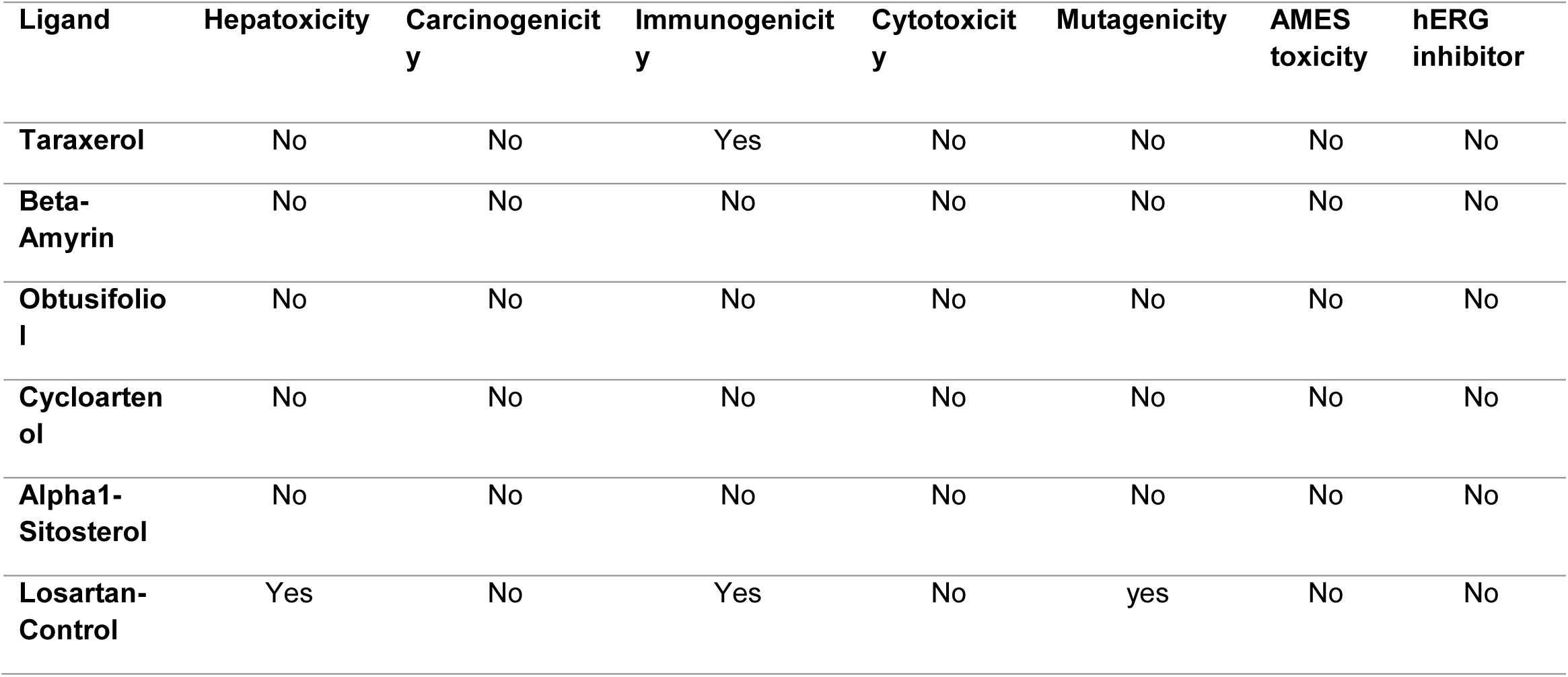
Toxicity profile of selected compounds.

Carcinogenicity and mutagenicity are increasingly significant among the various toxic categories because of their severe impacts on human healthTop of Form. The control compound Losartan (CID: 3961) had shown some carcinogenic and mutagenic activity. All the selected compounds showed safe toxicity profile. We wanted to determine if the pharmacological properties of the selected ligands correlated with each other. Therefore, we performed a Pearson correlation coefficient matrix analysis. Figure 3 shows a significant correlation between WLogP and Bioavailability, hydrogen bond donor, hydrogen bond acceptor and molecular weight with a p-value of <0.001. Additionally, the correlation between the molecular weight, WLopP and rotatable bond donors had a p-value of <0.01, indicating significance. Finally, E sol showed a significant correlation with rotatable bond and gastrointestinal absorption (%), with a p-value of <0.05. Our analysis suggests that hydrogen bond acceptors, donors, molecular weight and WLogP are crucial properties, as they exhibit significant correlations with other properties. For the other properties, the significance levels were >0.05, indicating insufficient data to demonstrate a correlation between them.

**Figure 3:**
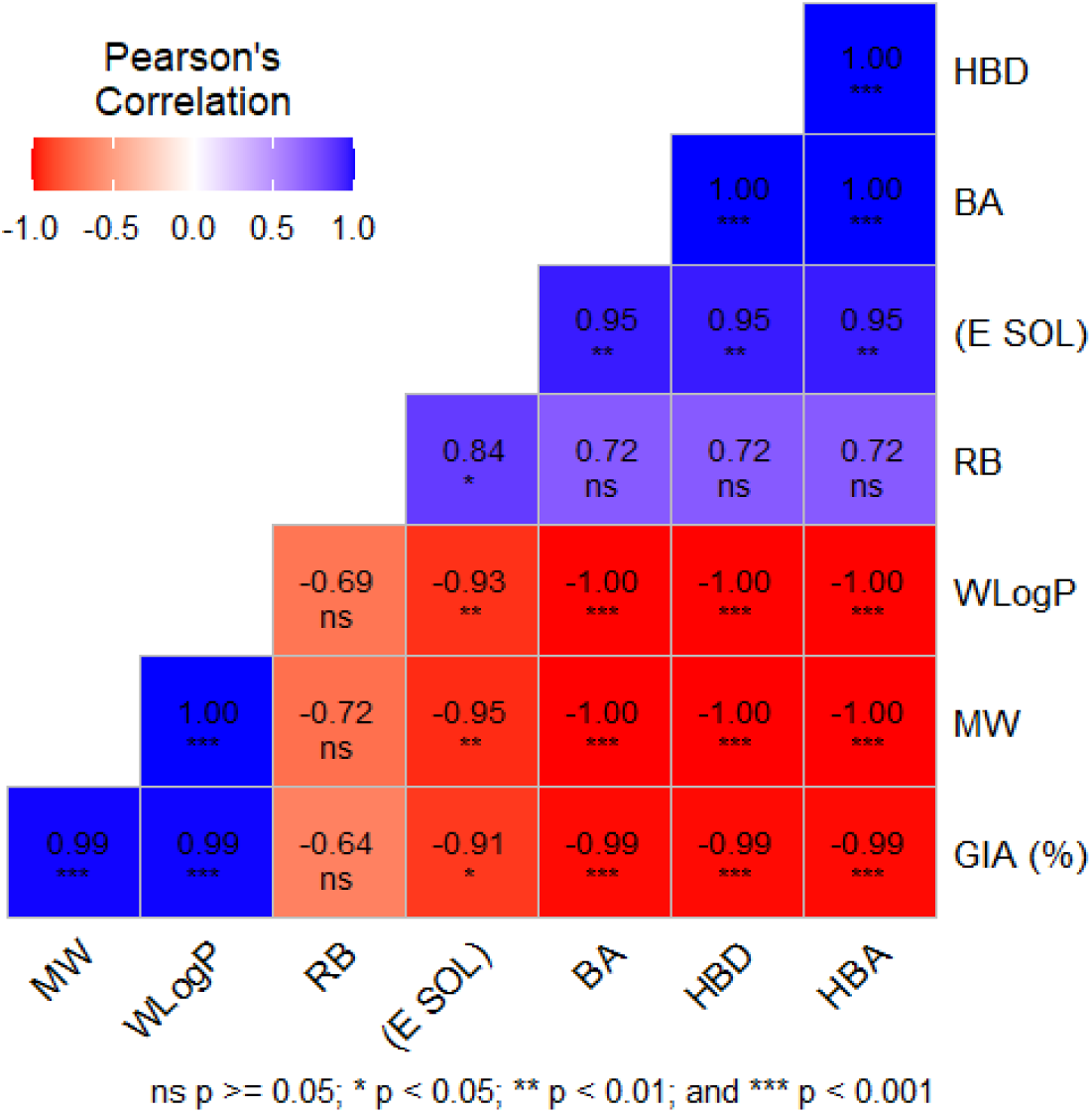
Analysis of Pearson correlation matrix showcasing the significance level between different pharmacological properties of the ligands.Here MW= molecular weight, HBD = hydrogen bond donor, HBA= hydrogen bond acceptor, RB= rotatable bonds, WLogP, (E SOL), GIA(%) =gastrointestinal absorption (%),BA= bioavailability.

### 3.6 Protein–ligand interactions

The intermolecular interactions of the top five protein-ligand complexes were examined. Both the hydrophobic interactions as well as the hydrogen bonds contribute to the shape and stability of the docked complexes. The protein-ligands interaction results and their bond types are summarized in Table1. The results are illustrated in Figure 4 which showcases 3D and 2D interactions between the AT1R and Phytocompounds.Taraxerol forms hydrogen bonds with amino acid residues Tyr35 and engages in 10 hydrophobic interactions. Beta-Amyrin, while lacking hydrogen bonds, exhibits six hydrophobic interactions with amino acid residues within the protein’s active site. Obtusifoliol and Cycloartenol form one and two hydrogen bonds, respectively, whereas Alpha1-Sitosterol shows none. The control, losartan, demonstrates a single hydrogen bond with residue Tyr35 and is involved in 11 hydrophobic interactions.

**Figure 4:**
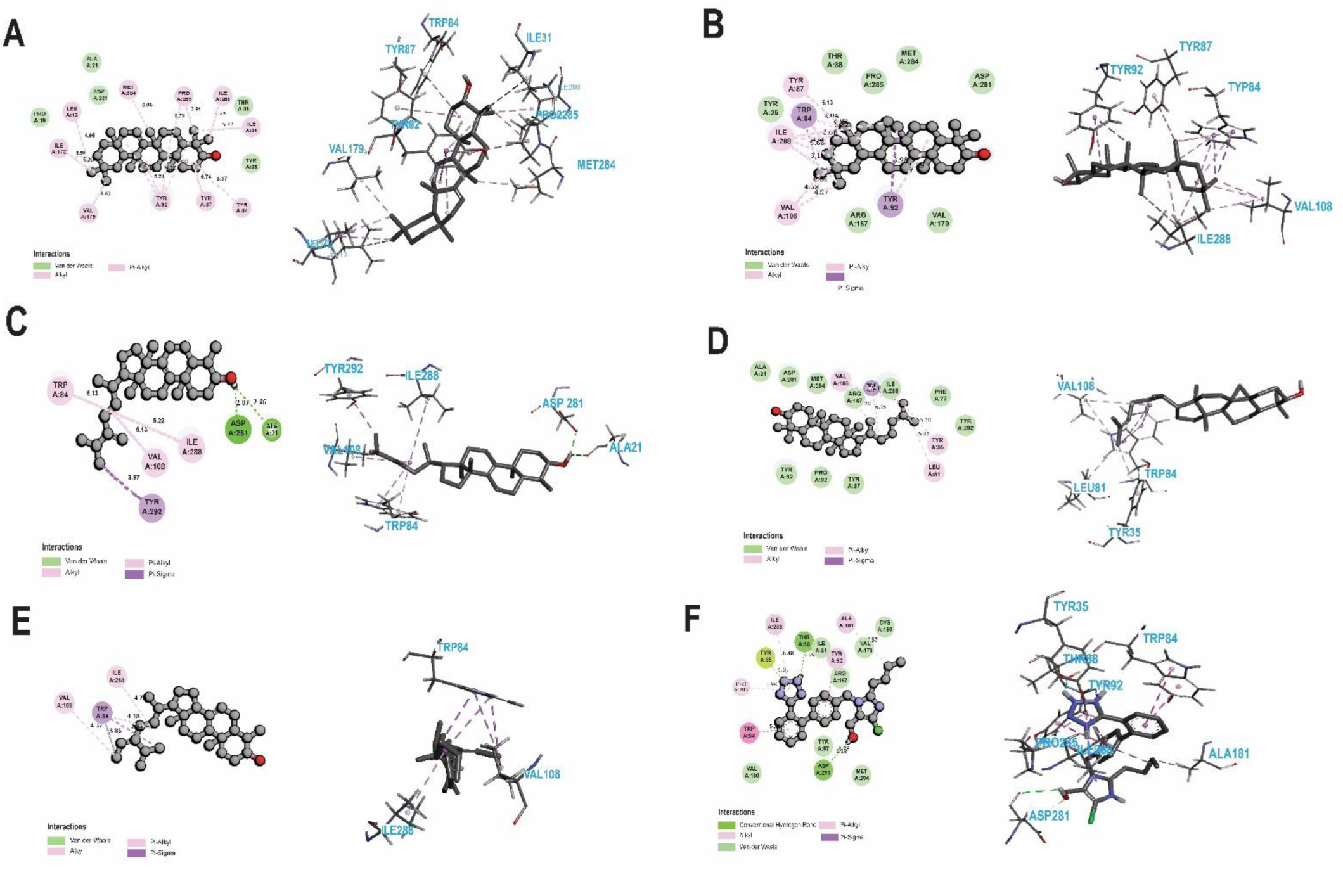
3D conformation (left) and 2D interaction plots (right) of best-docked complex molecules in the protein binding site for (A) Taraxerol - AT1R, B) Beta-Amyrin - AT1R (C) Obtusifoliol - AT1R, (D) Cycloartenol - AT1R (E) Alpha1-Sitosterol - AT1R (F) Control - AT1R

### 3.7 Molecular dynamic simulation

In computer-aided drug discovery, molecular dynamics simulation (MD simulation) is conducted to analyze the stability and intermolecular interactions within a protein-ligand complex in real-time. This method also enables the investigation of conformational changes within complex systems when they are subjected to simulated environments ^58^.

In this study, a 100 ns MD simulation of the protein bound to the specific ligand was performed to enhance comprehension of the protein’s conformational alterations within the complex. Initial analysis of intermolecular behavior was conducted utilizing the terminal snapshots from the 100 ns MDS trajectories

#### 3.7.1 RMSD analysis

In order to establish the stability profile of the chosen ligands, we performed RMSD analysis, a vital metric for evaluating the stability of compounds within the binding pocket of the receptor. In our RMSD analysis, we calculated the deviation in Cα atoms of the protein from its initial to final conformation during 100-ns molecular dynamics simulations. The stability of a conformation is determined by its deviation in simulations. A minimal deviation indicates substantial structural stability^59^. The Root Mean Square Deviation (RMSD) was calculated for two selected compounds, Taraxerol (CID-92097) and Beta-Amyrin (CID-73145), along with the control (CID-3961), as shown in (Figure 5).

**Figure 5:**
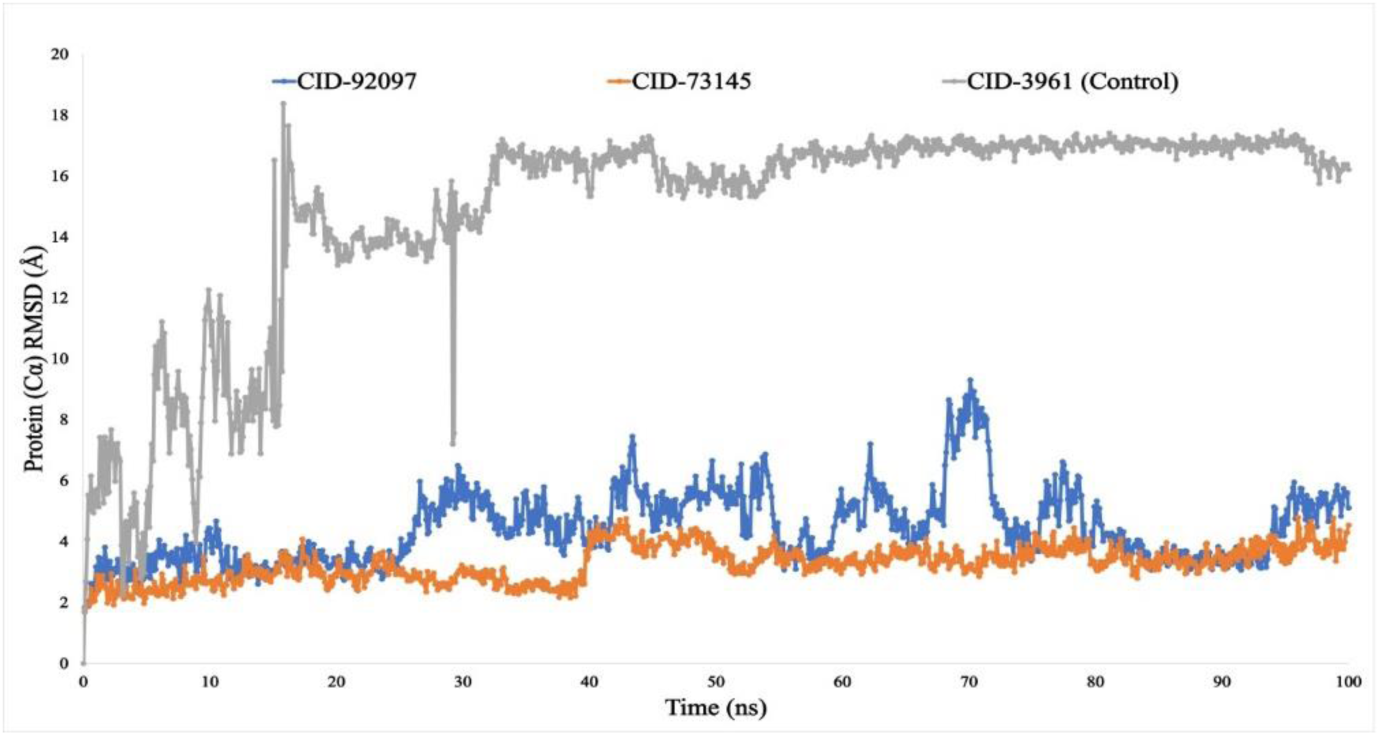
RMSD values extracted from C α of the protein–ligand docked complexes. The selected two ligands compound Taraxerol (CID-92097), Beta-Amyrin (CID-73145), and contol Losartan (CID-3961) in association with AT1R are depicted by blue, orange, and grey, respectively.

From the RMSD analysis the compound Losartan (Control) shows high fluctuation at the start of simulation point. within the 17–28 ns timeframe, remain stable but suddenly show higher fluctuation at 29ns. Towards the latter part of the simulation, from 30 to 100 ns, it demonstrates its stability. The Taraxerol (CID-92097) complex depicted by the blue line, it shows stability from the initial time point until the 26 ns mark in the simulation period. Subsequently, there is a slight increase in fluctuation observed from 26 to 80 ns. However, beyond 80 ns, the complex demonstrates a return to good stability, maintained up to 95 ns. The Beta-Amyrin (CID-73145) complex is depicted by the orange line, exhibiting stability from the initial time point until the 40 ns mark in the simulation period. Following this, there is a minor fluctuation observed from 41 to 55 ns. However, in the latter part of the simulation, specifically from 80 to 100 ns, it demonstrates its highest level of stability. Based on the preceding analysis, it is evident that among our selected ligands, Beta-Amyrin (CID-73145) complex exhibit minimum deviation, indicating the stability.

#### 3.7.2 RMSF analysis

Root Mean Square Fluctuation (RMSF) analysis can aid in characterizing and identifying local changes within the protein chain induced by specific ligand compounds interacting with particular residues. Elevated fluctuation scores imply increased flexibility and instability in the bonds, whereas lower scores suggest well-structured regions within a given protein-ligand complex^60^.

The RMSF values for the C-backbone atoms of Taraxerol (CID-92097), Beta-Amyrin (CID-73145), and losartan (CID-3961) in complex with the AT1R protein systems were calculated across a 100 ns trajectory to assess the impact of the attached ligand compounds on protein structural flexibility at specific residue positions, as shown in (Figure 6)

**Figure 6:**
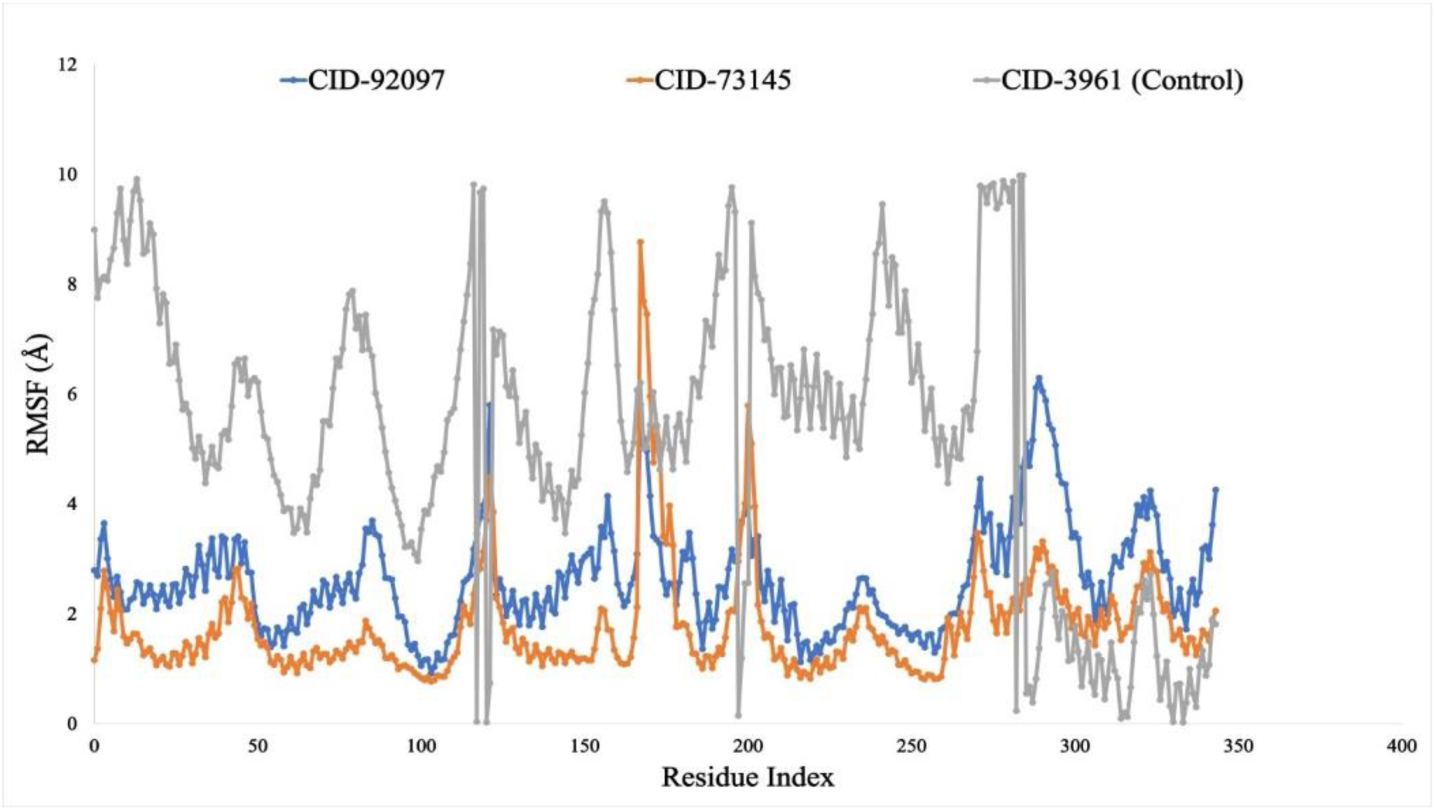
The RMSF values were calculated for the Cα atoms of the protein within the docked protein-ligand complexes. The selected two ligands compound Taraxerol (CID-92097), Beta-Amyrin (CID-73145), and control Losartan (CID-3961) in association with AT1R are depicted by blue, orange, and grey, respectively.

In the case of the control (grey line), many fluctuation peaks are observed entire the simulation time, that indicate the instability of control AT1R complex. For our selected ligands fluctuation observed at 121, 167, 200, 271, 289 and 323 no amino acid residues. The analysis of RMSF indicates that our ligands are forming stable complexes than our control drug.

### 3.8 Protein-ligand contact analysis

The MD simulation trajectories are employed for protein-ligand contact analysis, a pivotal step in the drug discovery process. This analysis offers valuable insights into atomic-level interactions between proteins and ligands. Throughout the 100 ns simulation duration, all compounds were observed to establish multiple interactions via hydrogen, hydrophobic, ionic, and Water Bridge bonding, which persisted until the conclusion of the simulation, contributing to the formation of a stable binding with the target protein (Figure 7). Taraxerol shows an interaction fraction value (IFV) of a maximum of 0.6 at the residue TYR 92. all portion of water bridges and the majority of hydrophobic bonds are involved in this interaction. Another compounds Beta–Amyrin shows of a maximum at 0.7 at the residue TYR 92 and hydrophobic bonds are involved. On the other hand, Losartan (control) shows the highest IFV among them is 1.1 for the residue PRO 285, where hydrogen, hydrophobic bonds and small portion water bridge bonds were involved

**Figure 7:**
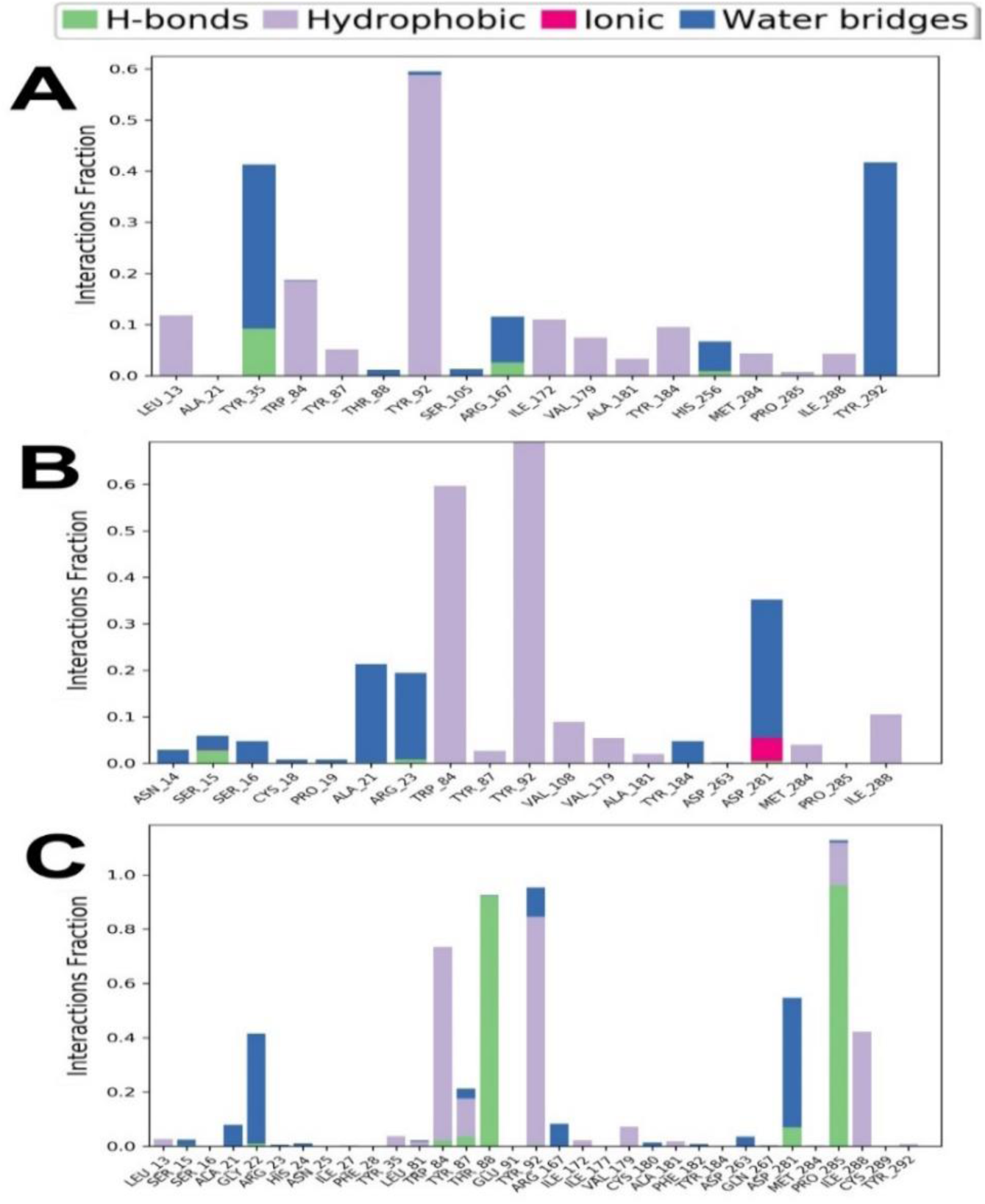
Protein-ligand interactions mapping for AT1R protein with (A) Taraxerol (CID-92097), (B) Beta-Amyrin (CID-73145), and (C) Losartan (CID-3961)

### 3.9 Post Simulation Binding Energy Analysis (MM/GBSA)

MMGBSA, or molecular mechanics-generalized Born surface area, is utilized to compute binding free energies and ligand strain energies for a collection of ligands interacting with a single receptor. A more negative value for the free energy of binding (MMGBSA_dG_Bind_vdW for complex-receptor-ligand) indicates a stronger binding affinity between the ligand compound and the targeted protein complex (Figure 8).

**Figure 8:**
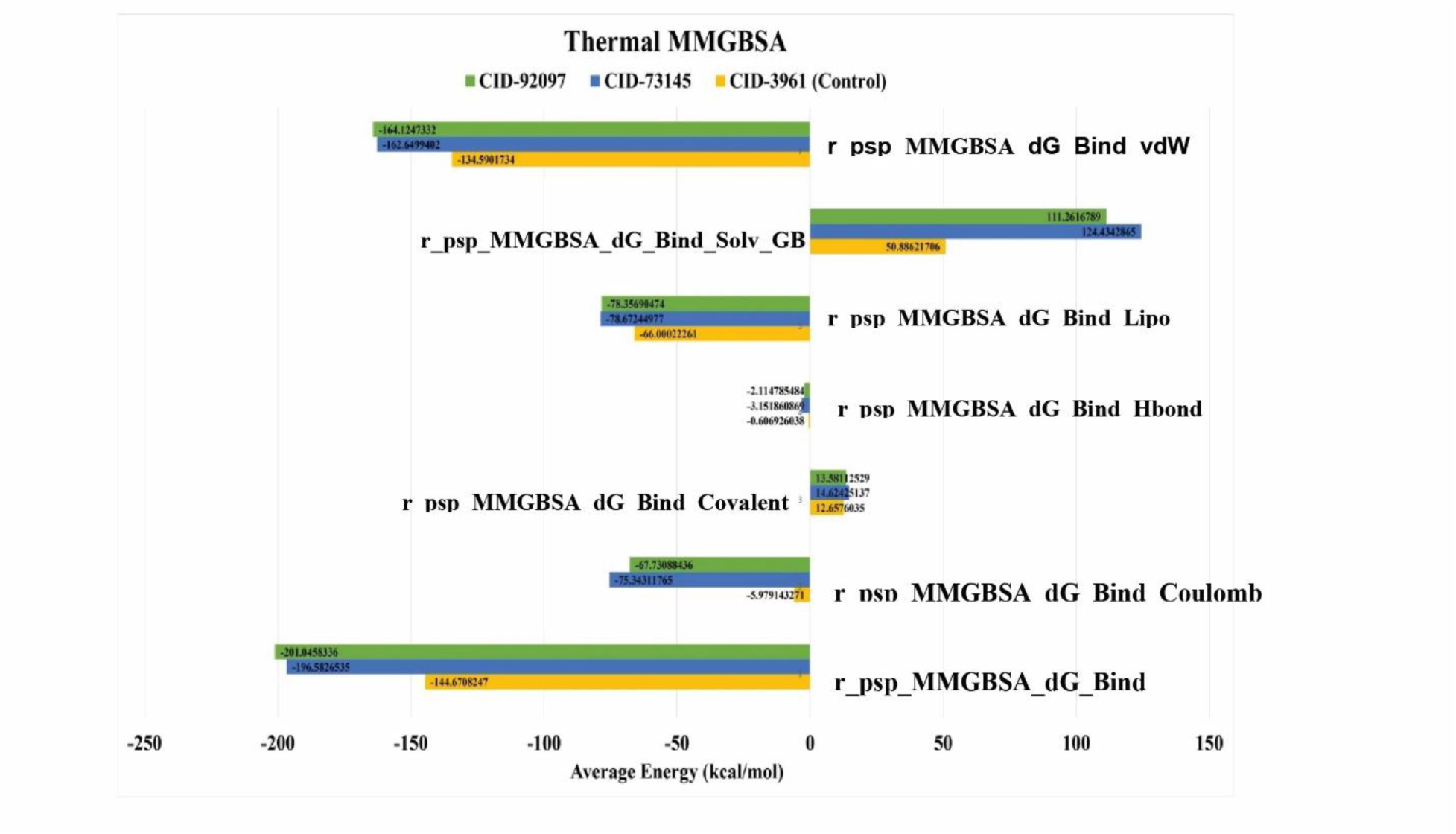
Post simulation trajectory MM/GBSA analysis of Taraxerol-green color, Beta Amyrin-green color, Lostartan (control)-yellow color with target protein AT1R complex (PDB ID: 4ZUD).

The results clearly illustrated that the Taraxerol and Beta Amyrin complexes with the receptor exhibited robust binding free energies of -164.124 kcal/mol and -162.649 kcal/mol, respectively. However, the compound losartan (Control drug) showed a lower binding free energy compared to our selected compound at -134.590 kcal/mol.

## 4. Discussion

AT1R is a critical factor of renin angiotensin-aldosterone system (RAAS) and is reported to be the only mediator of the biological actions of Angiotensin-II hormone ^9^. Ang-II is a vasoactive hormone, that maintains cardiovascular balance by regulating vascular resistance and blood volume. However, the extreme activation of AT1R by Ang II binding induces hypertension and facilitates to the outset of insulin-dependent diabetes. Impeding the activity of Ang-II, by inhibiting its binding with AT1R upon blood vessels leads to vasodilation, lowers blood pressure and potentially improves diabetes. Research indicates that AT1R antagonists have emerged as the preferred medication for individuals with hypertension, cardiovascular ailments, and type 2 diabetes due to their ability to regulate the excessive expression of AT1R ^61^. AT1 receptor blockers (ARBs) are those promising antagonists, can subtly reduce hypertension and diabetes via various mechanisms but lack in availability and possess adverse side effects. This meticulous study was employed to screen out novel drug-like candidate from *Nigella sativa* that could antagonize the Ang II.

CADD is an established method of exploring novel drugs via virtual screening from abundant candidates. It employs molecular docking to assess the most effective ligand that entails the interactions, binding activity and affinity between proteins and ligands. For the identification of our target protein inhibitor, we conducted multi-stage screening of 305 compounds of *Nigella sativa*. Before conducting molecular docking, the prediction of active sites of the protein were done. Initial molecular docking was performed and five compounds were considered for their promising binding affinities. The binding affinities of the five compounds Taraxerol, Beta-Amyrin, Obtusifoliol, Cycloartenol and Alpha1-Sitosterol are- -10.7, -10.7, -10.3, -10.3, -10.3 and -10.3 kcal/mol respectively, far greater than the control Losartan which is -8.2 kcal/mol. Lower binding affinity implies higher binding energy. The physiochemical, pharmacokinetic, and toxicity characteristics of the chosen potential drugs were implemented to ensure their safety, potency and efficacy after oral administration. Further, the interpretation of binding interaction the five compounds preserved required ligands within protein structures through essential weak intermolecular interactions like ionic, hydrogen, and hydrophobic bonding ^62^.

According to Lipinsk’s rule of 5 (RO5) compounds with two or fewer violations are deemed to have low toxicity, excellent absorption, and oral bioavailability. All examined ligands consistently meet these criteria and exhibit no infraction ^63^Top of Form. The poor intestinal absorption rate of a drug candidate is a drawback and makes it unproductive, irrespective of its high effective binding to the receptor ^64^. The absorption rate of our selected ligands was very satisfactory because the GIA percentage was very hing. Molecular weight and logP (logarithm of the partition coefficient) value are two considerable ADME properties, that significantly influence the biological membrane permeability of the drug molecule, lipophilicity and hydrophilicity. Higher molecular weight prevents the entrance of a drug molecule into the membrane barrier. A higher LogP value often indicates increased lipophilicity and potentially enhanced drug absorption, while a lower LogP value may suggest reduced efficacy due to insufficient membrane penetration. The lipophilicity of the selected 5 compounds was greater than the control (4.11) indicating that they can be better untaken by the AT1R membrane receptor. LogS, the logarithm of solubility, provides information about a substance’s hydrophilicity (water solubility) and is typically preferred to hold the lowest possible value ^55^. The LogS of these compounds lower than the control implying better solubility. The satisfactory number of H-bond donors and H-bond acceptors make a major impact in the drug’s interaction with its target, along with its other PK properties. The sufficient number of rotatable bonds is linked with oral bioavailability.

Assessing toxicity is a crucial aspect of drug development, as it helps identify potential adverse effects of a drug molecule in humanized environments, plants, or animals. Often clinical trials are employed on model animal or human to analyze the toxicity but the process is time-consuming and expensive and possess higher risk as well. Computational toxicity prediction involves no animal or human trials rather it is time-saving, cost-effective and provides a higher success rate in assessing toxicity profile for preclinical drug designing ^65^. The selected 5 compounds, evaluated in ProTox-3.0. and pkCSM web tool, hold negative toxicity profile showing no hepatoxicity, cardiotoxicity, carcinogenicity, immunogenicity and AMES toxicity. The selected compounds exhibited slightly immunogenic whereas other parameters presented favorable toxicity profile.

Molecular dynamics simulations play a crucial role in understanding the dynamic behavior of atoms within ligand-receptor complexes. We calculated the RMSD of the C-alpha backbone at the 100 ns timeframe to evaluate system stability and monitor structural changes over time. The distance-based RMSD values showed smaller magnitudes, indicating higher stability within the complexes. The Root Mean Square Deviation (RMSD) was analyzed for Taraxerol (CID-92097), Beta-Amyrin (CID-73145), and the control Losartan (CID-3961). Losartan initially showed high fluctuation, stabilized between 17–28 ns, fluctuated again at 29 ns, and then remained stable from 30 to 100 ns. The Taraxerol complex was stable until 26 ns, experienced slight fluctuations from 26 to 80 ns, and regained stability from 80 to 95 ns. Beta-Amyrin (CID-73145) demonstrated the least deviation, indicating the highest stability among the ligands tested. Root Mean Square Fluctuation (RMSF) analysis assesses local changes in protein chains due to ligand interactions, with high values indicating instability and low values suggesting well-structured regions. RMSF values for Taraxerol (CID-92097), Beta-Amyrin (CID-73145), and Losartan (CID-3961) in complex with the AT1R protein were calculated over a 100 ns trajectory. The control showed many peaks, indicating instability. Fluctuations for the selected ligands occurred at residues 121, 167, 200, 271, 289, and 323. RMSF analysis demonstrates that our ligands form more stable complexes than the control drug. Finally, we calculated the binding free energy, where the Taraxerol and Beta-Amyrin complexes showed strong binding energies of -164.124 kcal/mol and -162.649 kcal/mol, respectively, while the control drug Losartan had a lower binding energy of -134.590 kcal/mol.

Consequently, this investigation consolidated that the Beta-amyrin is the most potent candidate as ARB by employing numerous analyses including MDS. Beta-amyrin is a key triterpenoid bioactive compound, impacts blood sugar levels and lipid profiles. F.A. Santos et all demonstrated that Mice treated with β-amyrin exhibited a significant decrease in STZ-induced blood glucose (BG) levels and effectively lowered the elevated plasma glucose levels ^66^. This could lead to improved insulin sensitivity, boosting insulin production, and decrease in the blood-glucose that possibly reduce glucose absorption ^67^, Thus, Beta-amyrin act to be effective in diabetes for its antihyperglycemic and hypolipidemic effects ^66^. The other compound Taraxerol, is also a member of triterpenoid, previously proven to be a cardioprotectant, reducing the cardiotoxicity from clinical studies ^68^. Taraxerol has demonstrated various pharmacological effects, such as antitumor ^69^, antioxidant, antimicrobial ^70^, anti-inflammatory, antidiabetic ^71^, and anti-Alzheimer attributes ^72^.

We also wanted to find out whether there are any substantial health risks associated with these promising AT1R inhibitors. All of our finalized ligands don’t show any risks in laboratory-based studies. Although the ligands have slight problems in terms of bioavailability, and solubility, these issues can easily be solved during the drug development process, stages, and dosage formulation.

In this study, Beta-amyrin and Taraxerol manifests as novel and potent candidates for developing effective treatments for diabetic-hypertensive patient. Further in vitro and in vivo studies are strongly recommended to thoroughly evaluate the biochemical, preclinical, and clinical implications. The findings of this study will provide essential insight for the development of target-specific diabetic-hypertensive therapy.

## 5. Conclusion

In conclusion, over-activation of AT1R by the association of Ang II hormone leads to vasoconstriction, resulting in hypertension and delicately influencing T2DM. In the quest for novel phytochemical ARB drug candidates, we conducted a virtual screening of *Nigella sativa* due to its numerous therapeutic benefits, including diabetes, tumor, hypertension, and inflammation. Computer-based molecular docking was employed to anticipate potential bioactive compounds based on higher binding affinity, screening out five compounds showing desired pharmacokinetic properties, bioavailability, drug-likeness, efficacy, and potency as safe AT1R inhibitors. Subsequent investigations such as molecular dynamics simulation and MMGBSA facilitated the identification of both Beta-amyrin and Taraxerol as viable, efficacious, and non-toxic inhibitors of AT1R. This study highly represents the computer-aided investigation of compounds from *Nigella sativa* as an anti-hypertensive agent, confirming its biochemical and clinical advantages in targeting AT1R.

## Acknowledgement

The authors are grateful to Maksudul Islam for sharing his knowledge with us while conducting our research.

## Authors contributions

T.R.T and M.M designed, planned, and carried out the experiment. T.R.T conducted the research and also prepared this manuscript. M.M. participated in data analysis. S.K. and M.S.G guided the entire study and extensively revised this manuscript. M.N.H., P.B. and M.M.I.T. contributed to data processing and analysis. All authors were involved in the research and endorsed the final manuscript.

**Appendix A. Supplementary data**

## Abbreviations

AT1R: Angiotensin II Type1 Receptor
ARBs: AT1 receptor blockers
ADMET: Adsorption, Distribution, Metabolism, excretion and Toxicity
PDB: Protein Data Bank
RMSD: Root Mean Square Deviation
RMSF: Root Mean Square Fluctuation

