## Supporting Information Table S1 for "Computational screening of potential AT1R inhibitors from *Nigella sativa* for diabetic-hypertensive therapy"

**Supplementary Table- List of 305 Compounds extracted from *Nigella staiva* that were observed in this study.**

| No Compound id Compound name | Compound CID No | Docking score (kcal/mol) |
|---|---|---|
| Myristic acid | CID: 11005 | -5.8 |
| Myrtenol | CID: 10582 | -6.1 |
| (3S,4S)-4-ethenyl-4-methyl-3-prop-1-en-2-ylcyclohexene | CID: 10607083 | -6.5 |
| beta-Bisabolene | CID: 10104370 | -7.4 |
| Nigellidine | CID: 136828302 | -8.8 |
| O-Cymene | CID: 10703 | -6.3 |
| M-Cymene | CID: 10812 | -6.6 |
| Dillapiol | CID: 10231 | -6.2 |
| Thymoquinone | CID: 10281 | -6.6 |
| Carvacrol | CID: 10364 | -6.6 |
| Geranylacetone | CID: 1549778 | -6.3 |
| Thymol methyl ether | CID: 14104 | -6.3 |
| Nigellicine | CID: 11402337 | -5.6 |
| 2-Tridecanone | CID: 11622 | -5.7 |
| Pinocarvone | CID: 121719 | -6 |
| (-)-Butyrospermol | CID: 12302182 | -10.9 |
| Cycloeucalenol | CID: 101690 | -10.1 |
| Pentadecanoic acid | CID: 13849 | -5.7 |
| 2-(4-Methylphenyl)propan-2-ol | CID: 14529 | -6.4 |
| Lauric acid | CID: 3893 | -5.8 |
| Decanoic acid | CID: 2969 | -5.4 |
| Myristicin | CID: 4276 | -6.7 |
| Pimara-8(14),15-diene | CID: 440909 | -8.6 |
| Dithymoquinone | CID: 398941 | -8.8 |
| Myrcene | CID: 31253 | -5.9 |
| Coumarin | CID: 323 | -6.7 |
| Nonanal | CID: 31289 | -5.2 |
| Eugenol | CID: 3314 | -6.4 |
| Tetradecanal | CID: 31291 | -6.1 |
| 4-Isopropylbenzaldehyde | CID: 326 | -6.5 |
| alpha-Fenchene | CID: 28930 | -5.6 |
| (-)-Germacrene A | CID: 9548706 | -7.6 |
| 2,4-Decadienal | CID: 5283349 | -6 |
| (-)-beta-Bourbonene | CID: 62566 | -7.8 |
| gamma-Terpinene | CID: 7461 | -6.3 |
| (1S,2S,7S,8S)-2,6,6,9-tetramethyltricyclo[5.4.0.02,8]undec-9-ene | CID: 91753627 | -7.5 |
| Methyl linoleate | CID: 5284421 | -6.2 |

| | | |
|---|---|---|
| Longifolene | CID: 289151 | -7.4 |
| 2-Decenal | CID: 5283345 | -5.6 |
| Stearic acid | CID: 5281 | -5.6 |
| alpha-Ionone | CID: 5282108 | -6.7 |
| gamma-Himachalene | CID: 577062 | -8.1 |
| Longicyclene | CID: 564934 | -7.8 |
| p-Cymene | CID: 7463 | -6.3 |
| 2,2,5-Trimethyl-4-cyclohepten-1-one | CID: 88268 | -6.2 |
| Methyl geranate | CID: 5365910 | -6.6 |
| Cholesterol | CID: 5997 | -9.9 |
| 2-Pentadecanone | CID: 61303 | -6.5 |
| Myrtenal | CID: 61130 | -5.7 |
| Thymol | CID: 6989 | -6.8 |
| Obtusifoliol | CID: 65252 | -10.3 |
| Methyleugenol | CID: 7127 | -6.5 |
| Acetyleugenol | CID: 7136 | -6.8 |
| Estragole | CID: 8815 | -6.4 |
| 1-Decanol | CID: 8174 | -5.2 |
| Tricyclene | CID: 79035 | -5.6 |
| Methyl palmitate | CID: 8181 | -5.5 |
| Linalyl acetate | CID: 8294 | -5.9 |
| Dodecanal | CID: 8194 | -5.7 |
| Hederagenin | CID: 73299 | -9.7 |
| Umbellulon | CID: 91195 | -6.3 |
| Palmitic acid | CID: 985 | -5.9 |
| Citronellyl acetate | CID: 9017 | -6.3 |
| Thymohydroquinone | CID: 95779 | -6.3 |
| Spathulenol | CID: 92231 | -7.5 |
| 1-Methyl-4-(prop-1-en-2-yl)benzene | CID: 62385 | -6.4 |
| Citronellyl butyrate | CID: 8835 | -7.5 |
| Isolongifolene | CID: 11127402 | -7.5 |
| 6,7-Dimethoxy-1-methylisoquinoline | CID: 20725 | -6.7 |
| Lophenol | CID: 160482 | -10.1 |
| Tridecanoic acid | CID: 12530 | -5.9 |
| beta-Longipinene | CID: 25203064 | -7.7 |
| beta-Cyclocitral | CID: 9895 | -6.1 |
| 2,10-Epoxypinane | CID: 93046 | -5.6 |
| Cyclosativene | CID: 519960 | -7.6 |
| alpha,3-Dimethylstyrene | CID: 70759 | -6.4 |
| 2-(4-Methylphenyl)propan-1-ol | CID: 95376 | -6.7 |
| Isoterpinolene | CID: 102443 | -6.5 |
| Benzaldehyde | CID: 240 | -5.5 |
| Eucalyptol | CID: 2758 | -6 |
| beta-Elemene | CID: 6918391 | -7 |
| (-)-alpha-Himachalene | CID: 11830551 | -8.1 |

| | | |
|---|---|---|
| Nigellimine n-oxide | CID: 69131015 | -6.5 |
| Limonene oxide, cis-(-)- | CID: 6452061 | -6.1 |
| Thujopsene | CID: 442402 | -8.1 |
| p-Mentha-1,3,8-triene | CID: 176983 | -6.3 |
| trans-Sabinene hydrate acetate | CID: 6427504 | -6.7 |
| cis-Chrysanthenyl acetate | CID: 6431301 | -6.3 |
| 24-Methylenecycloartanol | CID: 94204 | -10 |
| Arachidic acid | CID: 10467 | -5.6 |
| 4-Carvomenthenol | CID: 11230 | -6.2 |
| 2-Undecanone | CID: 8163 | -6 |
| (1R)-2-methyl-5-propan-2-ylbicyclo[3.1.0]hex-2-ene | CID: 637518 | -6.1 |
| 4-Isopropylbenzyl alcohol | CID: 325 | -6.3 |
| Apiole | CID: 10659 | -6.1 |
| (2Z,6E)-Farnesyl acetate | CID: 1551480 | -6.3 |
| alpha-Spinasterol | CID: 5281331 | -10 |
| alpha-Selinene | CID: 10856614 | -7.9 |
| 7-epi-alpha-Eudesmol | CID: 12304196 | -7.5 |
| d-Borneol | CID: 6552009 | -5.7 |
| Terpinolene | CID: 11463 | -6.3 |
| alpha-Hederin | CID: 73296 | -10 |
| (E,Z)-farnesol | CID: 1549109 | -6.9 |
| Farnesol | CID: 445070 | -6.5 |
| Cycloartenol | CID: 92110 | -10.3 |
| alpha-Terpinene | CID: 7462 | -6.2 |
| Neryl acetate | CID: 1549025 | -6.1 |
| alpha1-Sitosterol | CID: 9548595 | -10.3 |
| beta-Farnesene | CID: 5281517 | -6.9 |
| Palmitoleic acid | CID: 445638 | -5.9 |
| Taraxerol | CID: 92097 | -10.7 |
| Zingiberene | CID: 92776 | -7 |
| Humulene | CID: 5281520 | -7.7 |
| Anethole | CID: 637563 | -6.4 |
| Farnesyl acetate | CID: 638500 | -6.9 |
| Citral | CID: 638011 | -5.8 |
| (+)-gamma-Cadinene | CID: 6432404 | -7.9 |
| Oleic acid | CID: 445639 | -6.2 |
| (Z)-gamma-bisabolene | CID: 3033866 | -6.9 |
| Tirucallol | CID: 101257 | -10.1 |
| Cinnamaldehyde | CID: 637511 | -5.9 |
| Pulegone | CID: 442495 | -6.4 |
| (-)-7-Epi-alpha-selinene | CID: 10726905 | -7.5 |
| (+)-delta-Cadinene | CID: 441005 | -7.9 |
| (-)-cis-Carveol | CID: 330573 | -6.4 |
| (-)-trans-Carveol | CID: 94221 | -6.2 |

| | | |
|---|---|---|
| Camphor | CID: 2537 | -5.7 |
| Linalool | CID: 6549 | -5.3 |
| alpha-Pinene | CID: 6654 | -5.7 |
| Carvone | CID: 7439 | -6.6 |
| Gamma-nonalactone | CID: 7710 | -5.7 |
| Isopropyl myristate | CID: 8042 | -5.3 |
| beta-Pinene | CID: 14896 | -5.9 |
| alpha-Terpineol | CID: 17100 | -6.1 |
| Sabinene | CID: 18818 | -6 |
| D-Limonene | CID: 440917 | -6.3 |
| Sabinene hydrate | CID: 62367 | -6.2 |
| beta-Amyrin | CID: 73145 | -10.7 |
| (+)-trans-Piperitenol | CID: 85568 | -6.1 |
| alpha-Curcumene | CID: 92139 | -7.3 |
| Campesterol | CID: 173183 | -9.9 |
| Epizonarene | CID: 595385 | -7.8 |
| Levomenol | CID: 442343 | -7.5 |
| Phytol | CID: 5280435 | -6.5 |
| Linolenic acid | CID: 5280934 | -6.4 |
| (1S,4E,9S)-4,11,11-trimethyl-8-methylidenebicyclo[7.2.0]undec-4-ene | CID: 6429301 | -7.4 |
| Isocaryophyllene | CID: 5281522 | -7.8 |
| (Z)-beta-Ocimene | CID: 5320250 | -5.5 |
| 24-Methylenelophenol | CID: 5283640 | -10.1 |
| gamma-Elemene | CID: 6432312 | -7.2 |
| alpha-Calacorene | CID: 12302243 | -9.1 |
| (-)-Carvomenthone | CID: 6432474 | -6.2 |
| 3,7-Dimethyloct-6-en-3-ol | CID: 86749 | -5.6 |
| Longiborneol acetate | CID: 91752502 | -7.3 |
| beta-Selinene | CID: 442393 | -7.2 |
| (E,Z)-2,4-Decadienal | CID: 6427087 | -5.5 |
| Caswell No. 264AB | CID: 442359 | -7.9 |
| alpha-Phellandrene | CID: 7460 | -6.5 |
| beta-Caryophyllene | CID: 5281515 | -7.7 |
| (E)-beta-ocimene | CID: 5281553 | -6.1 |
| beta-Sitosterol | CID: 222284 | -9.6 |
| Stigmasterol | CID: 5280794 | -10.2 |
| Bornyl acetate | CID: 93009 | -6.5 |
| Camphene | CID: 6616 | -5.9 |
| 2-Cyclohexen-1-ol, 2-methyl-5-(1-methylethenyl)-, acetate, cis- | CID: 102024 | -7.2 |
| cis-Pinocarveol | CID: 10931630 | -6 |
| 2-Cyclohexen-1-ol, 3-methyl-6-(1-methylethyl)-, (1R,6S)-rel- | CID: 85567 | -5.9 |
| cis-Sabinene hydrate | CID: 62367 | -6.2 |

| | | |
|---|---|---|
| Copaene | CID: 12303902 | -7.6 |
| Stigmastanol | CID: 241572 | -9.8 |
| 6-Epi-beta-bisabolol | CID: 12300148 | -7.5 |
| Bicyclo[2.2.1]heptan-2-ol, 1,7,7-trimethyl-, formate, (1R,2R,4R)-rel- | CID: 23623867 | -5.8 |
| Limonene | CID: 22311 | -6.4 |
| Linoleic acid | CID: 5280450 | -6.9 |
| (+)-trans-Limonene oxide | CID: 449290 | -6 |
| 2-Cyclohexen-1-ol, 1-methyl-4-(1-methylethyl)-, trans- | CID: 122484 | -6.3 |
| trans-Verbenol | CID: 89664 | -6 |
| alpha-Copaene | CID: 19725 | -7.7 |
| alpha-Fenchyl alcohol | CID: 439711 | -6 |
| 3-Buten-2-one, 4-(1,2,6,6-tetramethyl-2-cyclohexen-1-yl)- | CID: 5371122 | -6.8 |
| (1R,4S,5R)-4-methoxy-4-methyl-1-propan-2-ylbicyclo[3.1.0]hexane | CID: 101850207 | -5.9 |
| trans-Sabinene hydrate | CID: 12315151 | -6.2 |
| trans-4-Methoxythujane | CID: 71338689 | -6.1 |
| trans-4-Thujanol | CID: 20055523 | -6 |
| trans-alpha-Bergamotene | CID: 6429302 | -7.1 |
| gamma-Thujaplicin | CID: 12649 | -6.8 |
| Alloisolongifolene | CID: 1268122 | -7.3 |
| 3-Methylcatechol | CID: 340 | -6.1 |
| Hexadecyl butyrate | CID: 522033 | -5.8 |
| alpha-Santalyl acetate | CID: 6445771 | -6.9 |
| 2'-Hydroxy-5'-methoxyacetophenone | CID: 69714 | -6 |
| Hederagenin | CID: 73299 | -9.7 |
| Thiamine | CID: 1130 | -6.3 |
| Myristic acid | CID: 11005 | -6 |
| Heptadecanoic acid | CID: 10465 | -5.6 |
| Pyridoxine | CID: 1054 | -6 |
| Hexadecane | CID: 11006 | -4.9 |
| beta-Bisabolene | CID: 10104370 | -5.4 |
| Nigellidine | CID: 136828302 | -8.8 |
| Riboflavin | CID: 493570 | -7.4 |
| Thymoquinone | CID: 10281 | -6.5 |
| Carvacrol | CID: 10364 | -6.6 |
| Decane | CID: 15600 | -5.6 |
| p-Menth-3-en-1-ol | CID: 11468 | -6.1 |
| 2-Tridecanone | CID: 11622 | -5.5 |
| D-arabinonic acid | CID: 122045 | -5.2 |
| Pinocarvone | CID: 121719 | -5.9 |
| 2-(4-Methylphenyl)propan-2-ol | CID: 14529 | -6.5 |
| 2-(Hydroxymethyl)-6-[3-[3-[3,4,5- | | -4.5 |

| | | |
|---|---|---|
| trihydroxy-6-(hydroxymethyl)oxan-2-yl]oxypropoxy]propoxy]oxane-3,4,5-triol | CID: 4205683 | |
| Myristicin | CID: 4276 | -6.7 |
| Pimara-8(14),15-diene | CID: 440909 | -8.6 |
| Dithymoquinone | CID: 398941 | -7.9 |
| 4-Methoxybenzaldehyde | CID: 31244 | -5.6 |
| Myrcene | CID: 31253 | -5.9 |
| 4-Isopropylbenzaldehyde | CID: 326 | -6.5 |
| gamma-Terpinene | CID: 7461 | -6.2 |
| Davanone D | CID: 88556 | -7.1 |
| (1R,2R,7R,8R)-2,6,6,9-tetramethyltricyclo[5.4.0.02,8]undec-9-ene | CID: 92042758 | -7.7 |
| Arachidonic acid | CID: 444899 | -6.3 |
| Longifolene | CID: 289151 | -7.4 |
| Myristoleic acid | CID: 5281119 | -5.6 |
| 5-Dehydro-avenasterol | CID: 44263331 | -9.8 |
| Longicyclene | CID: 564934 | -7.3 |
| p-Cymene | CID: 7463 | -6.4 |
| 2,5-Dimethoxy-p-cymene | CID: 6427071 | -6.6 |
| Folic | CID: 135398658 | -9.3 |
| Ascorbic acid | CID: 54670067 | -5.8 |
| Thymol | CID: 6989 | -6.8 |
| 4-Acetyl-1,4-dimethyl-1-cyclohexene | CID: 65289 | -6 |
| Estragole | CID: CID: 8815 | -6.3 |
| Tricyclene | CID: 79035 | -5.6 |
| Nonane | CID: 8141 | -5.2 |
| Hederagenin | CID: 73299 | -9.7 |
| Palmitic acid | CID: 985 | -5.4 |
| Nicotinic acid | CID: 938 | -5.4 |
| Citronellyl acetate | CID: 9017 | -6.2 |
| Thymohydroquinone | CID: 95779 | -6.3 |
| 24-Ethyllophenol | CID: 541368 | -10 |
| Vitamin E | | |
| Pentadecane | CID: 12391 | -6 |
| Monogalactosyl diglyceride | | |
| 1-(11Z-icosenoyl)-2-(9Z,12Z-octadecadienoyl)-sn-glycero-3-phosphoethanolamine | CID: 102515444 | -6.4 |
| Cyclosativene | CID: 519960 | |
| Eucalyptol | CID: 2758 | -6 |
| Tetradecane | CID: 12389 | -6.1 |
| trans-Sabinene hydrate acetate | CID: 6427504 | -6.7 |
| 24-Methylenecycloartanol | CID: 94204 | -10.2 |

| | | |
|---|---|---|
| 4-Carvomenthenol | CID: 11230 | -6.2 |
| 2-Undecanone | CID: 8163 | -5.8 |
| beta-Eudesmol | CID: 91457 | -7.4 |
| delta7-Avenasterol | CID: 12795736 | -9.8 |
| (1R)-2-methyl-5-propan-2-ylbicyclo[3.1.0]hex-2-ene | CID: 637518 | -6.1 |
| Apiole | CID: 10659 | -6.3 |
| alpha-Spinasterol | CID: 5281331 | -10 |
| d-Borneol | CID: 6552009 | -5.7 |
| Terpinolene | CID: 11463 | -6.1 |
| Cycloartenol | CID: 92110 | -10.3 |
| alpha-Terpinene | CID: 7462 | -6.3 |
| Silibinin | CID: 31553 | -9.5 |
| alpha-Eudesmol | CID: 92762 | -7.7 |
| Anethole | CID: 637563 | -6.4 |
| Oleic acid | CID: 445639 | -6.7 |
| (+)-Dihydrocarvone | CID: 24473 | -6.2 |
| Betulinic acid | CID: 64971 | -9.4 |
| Camphor | CID: 2537 | -5.8 |
| Linalool | CID: 6549 | -5.6 |
| alpha-Pinene | CID: 6654 | -5.7 |
| Carvone | CID: 7439 | -6.6 |
| 1-Methyl-3-propylbenzene | CID: 14092 | -6.9 |
| beta-Pinene | CID: 14896 | -5.9 |
| alpha-Terpineol | CID: 17100 | -6 |
| Sabinene | CID: 18818 | -6 |
| D-Limonene | CID: 440917 | -6.3 |
| beta-Amyrin | CID: 73145 | -10.7 |
| Nerol | CID: 643820 | -5.7 |
| Linolenic acid | CID: 5280934 | -6.4 |
| Stigmasta-5,22-dien-3beta-yl alpha-D-glucopyranoside | CID: 101689889 | -6.4 |
| Pentyl Undecanoate | CID: 11594069 | -6 |
| methyl (14Z,16Z)-octadeca-14,16-dienoate | CID: 11630737 | -5.6 |
| pentyl (E)-hexadec-12-enoate | CID: 24873803 | -5.9 |
| methyl (15E,17E)-nonadeca-15,17-dienoate | CID: 24873804 | -5.8 |
| pentyl (E)-pentadec-11-enoate | CID: 24873805 | -5.9 |
| (-)-Carvomenthone | CID: 6432474 | -6.2 |
| 1-Ethyl-2,3-dimethylbenzene | CID: 13621 | -6.3 |
| Octyl isobutyrate | CID: 61024 | -5.8 |
| Fenchone | CID: 14525 | -6.2 |
| p-Mentha-1,5,8-triene | CID: 527424 | -6.3 |

| | | |
|---|---|---|
| (2E,4E)-3,7-dimethylocta-2,4,6-trienal | CID: 11126441 | -6.1 |
| 1,2-Dihydronaphthalen-2-one | CID: 173398 | -8.3 |
| Naphthalen-1(2h)-one | CID: 12446728 | -6.8 |
| (E,Z)-2,4-Decadienal | CID: 6427087 | -6 |
| alpha-Phellandrene | CID: 7460 | -6.5 |
| Aromadendrene | CID: 91354 | -7.5 |
| Astragalin | CID: 5282102 | -8.1 |
| beta-Caryophyllene | CID: 5281515 | -7.7 |
| beta-Sitosterol | CID: 222284 | -9.8 |
| Bornyl acetate | CID: 93009 | -6.5 |
| Camphene | CID: 6616 | -5.9 |
| cis-Sabinene hydrate | CID: 62367 | -6.2 |
| Limonene | CID: 22311 | -6.4 |
| Linoleic acid | CID: 5280450 | -5.4 |
| trans-Sabinene hydrate | CID: 12315151 | -6.2 |
